# Non-specificity of Pitstop 2 in clathrin-mediated endocytosis

**DOI:** 10.1101/002675

**Authors:** Anna K. Willox, Yasmina M. E. Sahraoui, Stephen J. Royle

**Affiliations:** Department of Cellular & Molecular Physiology, Institute of Translational Medicine, University of Liverpool, Liverpool, UK; Division of Biomedical Cell Biology, Warwick Medical School, Gibbet Hill Road, Coventry, UK

**Keywords:** clathrin, endocytosis, pitstops, small-molecule inhibitor

## Abstract

Small molecule inhibitors of clathrin-mediated endocytosis are highly desired for the dissection of membrane trafficking pathways in the lab and for potential use as anti-infectives in the clinic. One inhibition strategy is to prevent clathrin from contacting adaptor proteins so that clathrin-mediated endocytosis cannot occur. “Pitstop” compounds have been developed which block only one of the four functional interaction sites on the N-terminal domain of clathrin heavy chain. Despite this limitation, Pitstop 2 causes profound inhibition of clathrin-mediated endocytosis. In this study, we probed for non-specific activity of Pitstop 2 by examining its action in cells expressing clathrin heavy chain harbouring mutations in the N-terminal domain interaction sites. We conclude that the inhibition observed with this compound is due to non-specificity, i.e. it causes inhibition away from its proposed mode of action. We recommend that these compounds be used with caution in cells and that they should not be used to conclude anything of the function of clathrin’s N-terminal domain.

## Introduction

The ability to inhibit clathrin-mediated endocytosis (CME) using a small molecule is highly desirable. Not only would this be an extremely useful tool for cell biological studies, it would also have the potential to be used in the clinic as an anti-infective during viral or bacterial infection. Until recently, only crude methods for inhibiting CME existed and none of these acted with high specificity (von Kleist and Haucke, 2012). The recent description of Pitstop compounds as specific clathrin inhibitors held much promise (von Kleist et al., 2011). However, the specificity of these compounds and what conclusions can be drawn from their use is debated (Dutta and Donaldson, 2012; Lemmon and Traub, 2012).

A key step in CME is the engagement of adaptor proteins by clathrin (McMahon and Boucrot, 2011). The pitstop compounds were designed to inhibit this interaction and thus block CME specifically. The N-terminal domain (NTD) of clathrin heavy chain (CHC) is a seven-bladed beta propeller that has four binding sites (ter Haar et al., 1998; Willox and Royle, 2012). These sites are shown in Figure 1 and described in detail in Table 1. The first of these to be discovered was the groove between blades 1 and 2 which can bind “clathrin-box” motifs (LΦXΦ[DE]) (Goodman et al., 1997; Krupnick et al., 1997; ter Haar et al., 2000). Second, a site in the centre of the propeller binds “W-box” motifs (PWXXW) (Drake and Traub, 2001; Miele et al., 2004). Third, a groove between blades 4 and 5 accommodates a third type of motif that may be related to a clathrin-box ([LI][LI]GXL) (Kang et al., 2009). Fourth, mutational analysis identified a final site probably between blades 6 and 7, although the sequence requirement for peptide ligands to bind at this site is still unknown (Willox and Royle, 2012).

**Figure 1.**
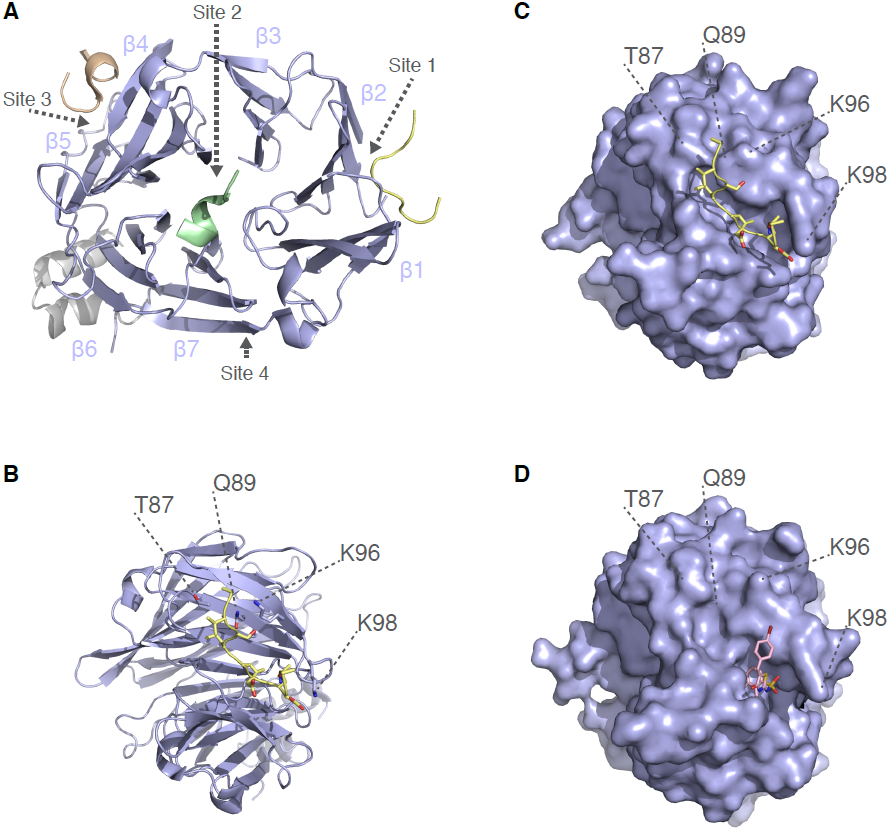
Structural view of interactions at the N-terminal domain of clathrin. A. Structural model of CHC(1-363), comprising the seven-bladed β-propeller (blue, 1-330) and linker (grey). The interaction sites listed in Table 1 are shown together with their respective ligands. The model is an alignment of three separate structures 1UTC, 1C9I and 3GD1. The blades of the β-propeller are numbered (β1-β7).
B. Cartoon view of the clathrin-box motif interaction site (Site 1). The ligand AVSLLDLA from β3-subunit of AP-3 (PDB code 1C9I) is shown together with residues that were targeted for the C+ mutation (see Table 1).
C. Surface view of the N-terminal domain in the same orientation as B, showing the clathrin-box motif peptide binding in the groove between blades 1 and 2. Positions of the residues targeted for mutation C+ are indicated.
D. Same view as C, but showing the NTD in complex with Pitstop 2 (PDB code 4G55). Pitstop 2 occupies the groove that clathrin-box motif-containing peptides bind, and this is believed to be its mode of action. No significant density for the ligand is found elsewhere on the NTD (von Kleist et al., 2011). Note that in 4G55, two alternative positions for side chains of Lys96 and Lys98 are present, conformation B is shown.

**Table 1.** Interaction sites on the N-terminal domain of clathrin heavy chain

|  | Adaptor Motif | Key residues | Examples | Mutant | Details |
| --- | --- | --- | --- | --- | --- |
| Site 1:<br>Clathrin-box motif | LØXØ[DE] | Ile80<br>Thr87<br>Gln89<br>Phe91<br>Lys96<br>Lys98 | AP-2 β2-subunit<br>Amphiphysin<br>β-arrestin 1 | C+ | T87A<br>Q89A<br>K96E<br>K98E |
| Site 2:<br>W-box motif | PWXXW | Phe27<br>Gln152<br>Ile154<br>Ile170 | Amphiphysin<br>SNX9 | D | Q152L<br>I154Q |
| Site 3:<br>β-arrestin 1L site | [LI][LI]GXL | Arg188<br>Gln192 | β-arrestin 1L<br>AP-2 β2-subunit | E | R188A<br>Q192A |
| Site 4:<br>Fourth site | Unknown | Glu11 | Unknown | G | E11K |

The pitstop compounds were designed to target only one of these sites: the clathrin-box motif site (von Kleist et al., 2011). Figure 1B shows the interaction of clathrin-box motifs with key residues in the groove between blades 1 and 2 of the NTD. Structural studies showed that Pitstop 1 and Pitstop 2 compounds occupy the same site on the NTD, and thus block the interaction of clathrin-box motif-containing peptides with the NTD of CHC *in vitro* (Fig 1C) (von Kleist et al., 2011).

Previously it was shown that any one of the four interaction sites on the CHC NTD is sufficient to support CME in human cells. Moreover, inhibition comparable to removal of the NTD, only occurs after all four sites have been mutated (Willox and Royle, 2012). Thus it is very surprising that pitstops, compounds that bind only at the CBM site on the NTD *in vitro*, can block CME effectively in cells (von Kleist et al., 2011). This raised questions about their specificity of action in CME (Lemmon and Traub, 2012).

Recent work has shown that Pitstop 2 inhibits clathrin-independent endocytosis in addition to blocking CME. The authors showed that clathrin-independent endocytosis of two different cargo proteins was inhibited by Pitstop 2 (Dutta et al., 2012), although whether the internalization of one of these cargos is truly clathrin-independent has been questioned (Stahlschmidt et al., 2014). This argues that the compound may either have additional off-target effects or that the inhibition of CME and clathrin-independent endocytosis is via a common mechanism that does not involve clathrin at all (Dutta and Donaldson, 2012; Dutta et al., 2012; Lemmon and Traub, 2012).

In this paper we set out to test the specificity of action of pitstop compounds in CME. Our goal was to establish what conclusions, if any, can be drawn about clathrin function using these compounds. In their critique of pitstops, Lemmon and Traub stated “it will certainly be interesting to determine whether pitstops interfere with clathrin-mediated endocytosis when only the surface chemistry of the clathrin-box site is altered by site-directed mutagenesis” (Lemmon and Traub, 2012). This is precisely the method of testing that we report here. Our results indicate that Pitstop 2 has inhibitory effects outside of the NTD of CHC. We suggest that results from experiments using pitstops should be treated with caution and these compounds should not be used to conclude anything about the function of the NTD of clathrin in cells.

## Materials and Methods

All cell culture reagents were from Life Technologies (UK) or Sigma Aldrich (UK), unless stated otherwise. Pitstop 1 and Pitstop 2 were gifts from Megan Chircop and Phil Robinson (Children’s Medical Research Institute, Sydney, Australia). All plasmids were available from previous work (Hood et al., 2013; Willox and Royle, 2012), with the exception of plasmids for expression of mutant MBP-CHC(1-1074)-His6 in bacteria. These were made by standard site-directed mutagenesis. All plasmids were verified by automated sequencing.

Cell culture and flow cytometry was as described previously (Willox and Royle, 2012). Briefly, HeLa cells were maintained in DMEM supplemented with 10% fetal bovine serum and 100 units/ml penicillin/streptomycin at 37 °C and 5% CO_2_. Cells were transfected using Lipofectamine 2000 (siRNA) or Gene Juice (plasmid, Merck) according to the manufacturers’ recommendations. We used a “two-hit” protocol: cells were transfected with siRNA on day 1, split and transferred to new plates on day 2 and then transfected with the appropriate pBrain plasmids on day 3. Six hours after transfection, medium was replaced with fresh growth medium containing 5 mM sodium butyrate to increase efficiency of transfection and expression. After 16 hours (day 4), the media was replaced and cells were assayed on day 5. The siRNA was directed against human CHC or a GL2 control siRNA was used followed by pBrain-GFP-CHC1 transfection.

Transferrin uptake was assayed using flow cytometry. Cells were first incubated for 30 minutes at 37°C in serum-free DMEM, with Pitstop compounds (30 *µ*M) or with DMSO control (0.1%). Cells were then trypsinized for 3 min at 37 °C, briefly pelleted and then incubated with 50 *µ*g/ml Alexa Fluor 647–conjugated transferrin for 5 min at 4°C. The cells were moved to 37°C and incubated for 10 minutes. Cells were pelleted, washed once in PBS, acid-washed twice (0.1 M glycine, 150 mM NaCl, pH 3), and resuspended in PBS containing 1% BSA.

Flow cytometry was done using a two-laser, four-color FACSCalibur flow cytometer (Becton Dickinson, Oxford, United Kingdom). Typically, 1000-10000 cells were analyzed per experiment. Cells were gated so that only cells expressing GFP above a threshold, identical for all constructs, were included in the analysis (as shown in Figure 2). The median Alexa Fluor 647-transferrin signal from the gated cells was used to compare and plot data..

**Figure 2.**
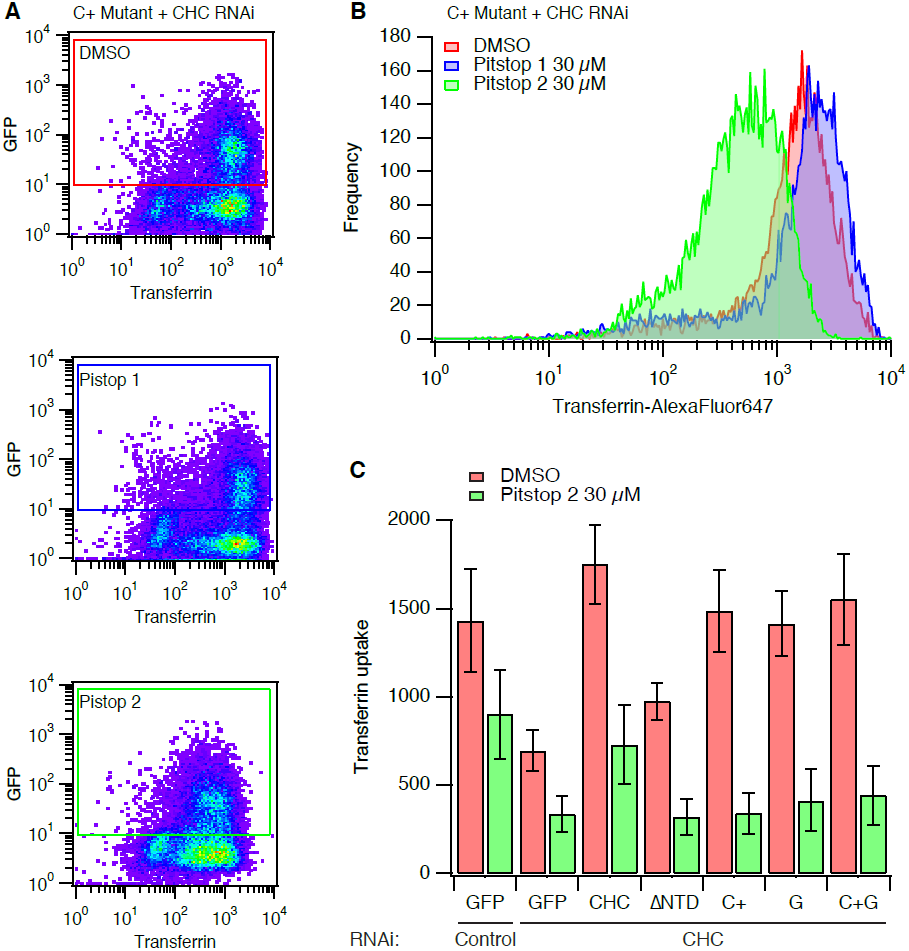
Pitstop 2 inhibits clathrin-mediated endocytosis non-specifically. A. Typical data from transferrin uptake experiments with clathrin-depleted cells expressing GFP-CHC(1-1675) (C+ mutant). Cells were treated with DMSO or Pitstop 1 (30 *µ*M) or Pitstop 2 (30 *µ*M) for 30 min during serum starvation, as indicated. A pseudocoloured bivariate histogram is shown for GFP intensity *versus* transferrin-Alexa647 fluorescence. The GFP-positive cells were a clearly discerned population that could be gated and analysed as indicated.
B. Histogram to show the frequency of cells with a given transferrin uptake in clathrin-depleted expressing full-length RNAi-resistant GFP-tagged CHC harbouring the C+ mutations. Histograms were generated from data gated as indicated in (A). Note the logarithmic scale on the x-axis.
C. Bar chart to summarise transferrin uptake experiments. Cells were transfected as indicated. Median transferrin-Alexa647 fluorescence values from cells gated for GFP expression are shown as mean ± s.e.m. from four independent experiments.

Protein purification and binding experiments were carried out as previously described (Hood et al., 2013). For *in vitro* binding assays involving TACC3, 50 *µ*g of GST or GST-tagged TACC3 was incubated with 2 *µ*g/ml Aurora A kinase (Millipore) or BSA, 2 *µ*g/ml GST-TPX2(1-43) and 10 mM MgATP for 2 hours at 30°C in reaction buffer (50 mM Tris.Cl, pH 7.5, 150 mM NaCl, 0.1 mM EGTA). This phosphorylated protein was then used for the bind reaction. For GST or GST-β2 appendage and hinge (616-951), proteins were not phosphorylated. For binding, GST-protein was incubated with 30 *µ*l of glutathione sepharose 4B in a total volume of 200 *µ*l NET-2 buffer (50 mM Tris.Cl, pH 7.5, 150 mM NaCl, 0.5% NP-40 substitute, containing 0.1 mg/ml of MBP-CHC(1-1074) or mutant versions. Proteins were incubated overnight with rotation at 4°C, then spun at 10,000 × g for 2 min. Supernatant was retained and beads were washed 4 times with 1 ml NET-2. 30*µ*1 of 2 X Laemmli buffer was added to the beads, they were denatured at 100°C for 5 min and half was analyzed by western blot along with 5 *µ*l of the supernatant.

Data analysis and presentation was done using Igor Pro 6.34 (Wavemetrics) or PyMol (DeLano Scientific). Figures were assembled in Adobe Illustrator CS5.1.

## Results

We have previously used a strategy to test the function of various CHC mutants by depleting endogenous CHC by RNAi and simultaneously expressing an RNAi-refractory version of CHC that is tagged with GFP (Willox and Royle, 2012). In the present study we again used this system and measured the uptake of fluorescent transferrin using flow cytometry. The uptake of transferrin is used because it is known to be by AP2-dependent CME (Motley et al., 2003). Using flow cytometry, the cells depleted of endogenous clathrin and expressing GFP-tagged proteins can be gated according to GFP fluorescence (Fig 2A) and the uptake within the gate analysed (Fig 2B). As previously described, transferrin uptake was inhibited by depletion of CHC and this inhibition was rescued by the expression of full-length CHC, but not by a CHC mutant lacking the N-terminal domain (ΔNTD) (Fig 2C). Three further CHC mutants were tested in parallel. These were: mutant C+ targeting the clathrin-box motif site, mutant G targeting the “fourth site” and mutant C+G which combined these two sets of mutations (Table 1). As described previously, all three CHC mutants could support CME to the same extent as wild-type CHC (Fig 2C).

In order to test the specificity of Pitstop 2, cells were pre-incubated with the compound (30 *µ*M) for 30 min during serum starvation. This treatment inhibited transferrin uptake in all conditions compared to DMSO control cells (Fig 2C). The amount of transferrin uptake in Pitstop 2-treated control RNAi cells was equivalent to that in control-treated CHC RNAi cells. In clathrin-depleted cells expressing either GFP or CHC mutants, Pitstop 2 caused a further inhibition of residual transferrin uptake (Fig 2C). The inhibition of transferrin uptake in CHC mutants C+ and C+G is particularly noteworthy as these mutants are predicted to be unable to bind peptides bearing clathrin-box motifs (Fig 1 and Table 1). Indeed, this is the rationale for the design of pitstop compounds. This result indicates a non-specific inhibitory action of Pitstop 2 on CME.

The inhibition of CME by Pitstop 2 was not a result of the drug acting on a subset of cells, but instead the inhibition was global (Fig 2A, B). Histograms of transferrin uptake in clathrin-depleted cells expressing the C+ mutant clearly show that the entire population of cells is shifted to the left, with a single peak (Fig 2B). These plots also shows that Pitstop 1, a compound that occupies the same NTD site, has no discernable effect (Fig 2A, B).

In a further iteration of this experiment we tested additional CHC mutants and uptake in every case was inhibited by pre-incubation with Pitstop 2 (Fig 3). These CHC mutants included mutant D and mutant E in which the W-box site or the β-arrestin 1L site were targeted by mutation, respectively (Table 1). In addition, a triple site mutant C+DE was also tested, this mutant has only one functional site, the fourth site (Willox and Royle, 2012). Again, CME in clathrin-depleted cells expressing these mutants was blocked by Pitstop 2 (Fig 3).

**Figure 3.**
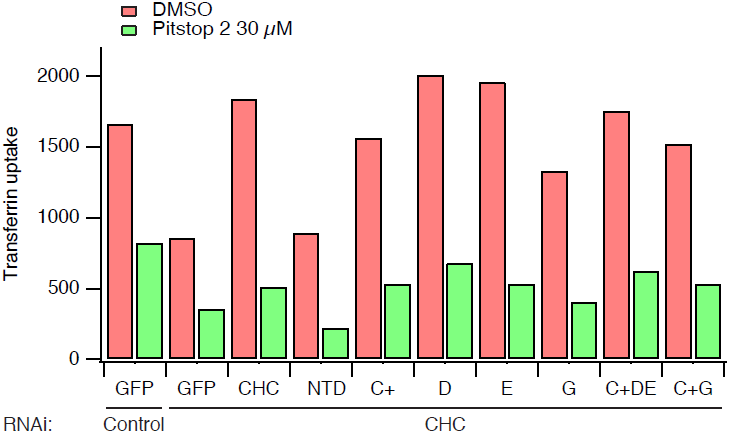
The inhibition of CME by Pitstop 2 is not confined to the clathrin-box site. Bar chart of a single transferrin uptake experiment. Cells were transfected as indicated. Median transferrin-Alexa647 fluorescence values from cells gated for GFP expression are shown.

Together, these observations suggest that Pitstop 2 acts non-specifically to inhibit CME. First, it is extremely unlikely that the mode of action of Pitstop 2 is as described: by solely blocking the clathrin-box motif-binding groove on the NTD. Second, the compound cannot discriminate between CHCs that harbour mutation of any of the interaction sites on the NTD.

The conclusion that Pitstop 2 acts non-specifically, that is, by not acting solely at the CBM-binding site rests on the C+ mutations being sufficient to disrupt the binding of clathrin-box motifs to the NTD of CHC. If they were not, it could be that there was residual binding to the mutant that is then inhibited by pitstop occupancy. To test this possibility, we analysed the *in vitro* binding of CHC fragments containing the NTD to GST-tagged recombinant β2 hinge and appendage domains (Fig 4A). MBP-tagged CHC(1-1074) bound tightly to GST-β2(616-951) but not to GST alone. Mutation of the CBM-binding site (mutant C+) or the β-arrestin 1L site (mutant E) prevented binding, while mutation of the other two sites individually had no effect. The C+ or E mutations did not cause gross changes in the protein structure, as binding to phosphoTACC3 remained intact (Fig 4B). These experiments indicate that C+ mutations are sufficient to prevent *in vitro* interactions with endocytic adaptors via this groove. The inhibition of CME by Pitstop 2 in clathrin-depleted cells expressing the C+ mutant CHC is therefore due to a non-specific activity.

**Figure 4.**
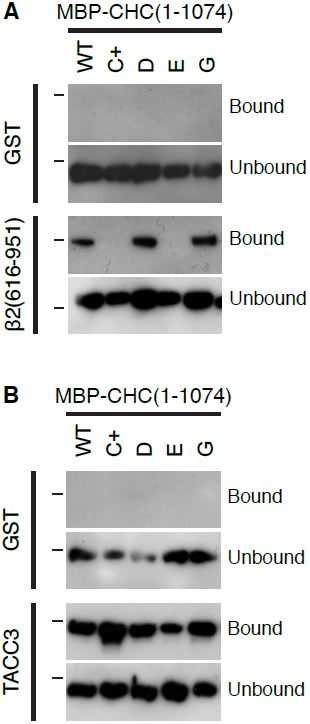
Mutation of the clathrin box site is sufficient to block interaction with clathrin box motif-containing proteins. Results of a typical binding experiment to analyse the role of the four interaction sites on the NTD of CHC. Wild-type MBP-CHC(1-1074)-His6 or indicated mutants were incubated with (**A**) GST or GST-β2 adaptin (616–951) or (**B**) GST or GST-TACC3-His6 that were phosphorylated with Aurora A/TPX2, before pull-down on glutathione beads. Bound, and a sample of unbound, material was separated by SDS-PAGE and transferred to nitrocellulose for western blotting using an anti-CHC (TD.1) antibody. Markers indicate position of 150 kDa.

## Discussion

Specific inhibitors of CME are highly desired for molecular dissection of trafficking pathways in the lab or for their potential use as anti-infectives in the clinic. Classical methods for inhibition of CME include treatment of cells with hypertonic sucrose or depletion of intracellular potassium (Heuser and Anderson, 1989; Larkin et al., 1983). While these methods are effective, they clearly have off-target effects which limit their use and so the design of more selective inhibitors is highly desired (Dutta and Donaldson, 2012).

At the turn of the last century, the model for how clathrin engages with the adaptor layer during CME was that the NTD of CHC binds AP-2 via a CBM in the linker region of β2 subunit of the heterotetrameric adaptor complex, AP-2 (Goodman et al., 1997; Krupnick et al., 1997; ter Haar et al., 2000). This rationale was used for the development of clathrin inhibitors, pitstops (von Kleist et al., 2011). However, multiple lines of evidence demonstrate that this original model was oversimplified. First, three other interaction sites on the NTD of CHC were subsequently identified (Drake and Traub, 2001; Kang et al., 2009; Miele et al., 2004; Willox and Royle, 2012). Second, the demonstration that multiple sites are used for clathrin function in yeast and human cells (Collette et al., 2009; Willox and Royle, 2012). Third, interaction sites on the leg of CHC may play a role in adaptor engagement (Edeling et al., 2006; Knuehl et al., 2006). Much of this information was known when the pitstops were first described. The effectiveness of Pitstop 2 in inhibiting CME directly contradicts the claims of its specific action. Lemmon and Traub explored this contradiction in their critique of pitstops (Lemmon and Traub, 2012).

In this short study we tested the extent of Pitstop 2 non-specificity. We found that Pitstop 2 inhibited CME regardless of which CHC mutant actually mediated endocytosis. Clathrin-depleted cells that expressed CHC with the C+ mutation, which blocks the binding of clathrin-box motifs to the NTD, can support normal CME. Pitstop 2 efficiently inhibited CME in these cells. Since CME in these cells is independent of interactions at the groove where Pitstop 2 is proposed to act, then the inhibitory action is a sign of non-specificity.

There is very little evidence that in cells, Pitstop 2 acts in the way it is does *in vitro*, i.e. binds in the groove that accommodates clathrin-box motif peptides. Clathrin and TACC3 remain on the mitotic spindle when cells are treated with Pitstop 2 at doses that inhibit CME (Smith et al., 2013). Since the C+ mutation in CHC prevents clathrin binding to the mitotic spindle in concert with phosphoTACC3 (Hood et al., 2013), this argues that the compound either does not bind at this site in cells or that its binding is too low affinity to outcompete the interaction of clathrin–TACC3 with microtubules.

What is the extent of the non-specificity of action? There are three realms of non-specific action that can be considered. The first is that Pitstop 2 acts non-specifically to block all four interaction sites on the NTD of CHC. We found that Pitstop 2 was inhibitory for all CHC mutants, including the C+DE mutant, which only has one operational site (Table 1). Since any of the four interaction sites can support CME (Willox and Royle, 2012), then this realm of non-specific action would explain our observations. The second is that it blocks other interaction sites on CHC outside of the NTD. This is difficult to test in cells because a CHC mutant that lacks all four interaction sites (ΔNTD or CDEG, in (Willox and Royle, 2012)) is unable to support endocytosis and so we cannot test for a non-specific inhibitory action of Pitstop 2 using this mutant. It is possible that the compound blocks other sites on CHC such as those on the ankle, but there is no experimental evidence to suggest that this is the case. The third realm is that the inhibitory action of Pitstop 2 is completely non-specific, i.e. not acting on CHC at all. *In vitro* binding assays showed that other interactions in membrane trafficking such as between amphiphysin 1 and α-subunit of AP-2 were only minimally affected by Pitstop 2, although this was not tested in cells (von Kleist et al., 2011). This argues against a scenario where pitstops non-specifically inhibit each and every peptide-peptide interaction in CME. However, the inhibition of clathrin-independent endocytosis indicates that Pitstop 2 acts on other non-clathrin target(s) (Dutta et al., 2012). The observation of changes in mitotic progression that may be unrelated to clathrin also point to off-target effects of this compound (Smith et al., 2013).

The next generation of 1,8-Naphthalimide clathrin inhibitors have now been reported (Macgregor et al., 2014). As a class of compounds, several inhibit NTD interactions *in vitro* yet only Pitstop 2 inhibits CME in cells (Macgregor et al., 2014). Pitstop 1 is apparently able to inhibit CME when microinjected into cells and it is assumed that other similar compounds cannot permeate cells to exert an inhibitory effect. The permeability of these compounds has not been tested directly and so it is possible that these compounds can enter cells and are selective for the clathrin-box motif site on clathrin. Like the C+ mutation this would not be sufficient to block CME, giving the impression that they are unable to enter cells. If this is the case then the inhibition of CME by Pitstop 2 is unusual for this class of compounds and would suggest an entirely non-specific action.

Our study highlights the non-selectivity of Pitstop 2 and in light of these concerns, we recommend that these compounds should not be used to draw conclusions about CHC NTD function in cells. Drugging protein-protein interactions is extremely challenging and any attempts to generate new clathrin inhibitors are much appreciated by cell biologists. Our understanding of how clathrin interacts with the adaptor layer is incomplete. Which of the four sites on the NTD are used and when? Is the ankle site essential for interactions? Answers to these questions may allow for further inhibitor design or rule out clathrin-adaptor interactions as a reasonable target for inhibition. We hope this critical appraisal of pitstop selectivity does not hinder further development of clathrin inhibitors, which would be a tremendous asset to the cell biologists toolbox. Hopefully in the future we will have truly selective inhibitors of key interactions within CME. Unfortunately, we are not at that point yet.

## Acknowledgements

We thank Samantha Williams for help with binding experiments, Richard Bayliss for advice on interpretation of PDB files, Saty Kaur and Laura Wood for critically reading the manuscript and other Royle Lab members for discussion. We are very grateful to Megan Chircop and Phil Robinson who generously donated the pitstop compounds that were used in this study. Thanks also to James Woodgett who encouraged SJR in 140 characters to write up this work.

### Competing interests

The authors have no competing interests to declare

### Author Contributions

AKW performed the flow cytometry experiments. YS did the binding experiments. SJR wrote the paper.

### Funding

This work was supported by a project grant (WT084569MA) from the Wellcome Trust and a Summer Studentship from the Wellcome Trust to YS.

## References

Collette, J. R., Chi, R. J., Boettner, D. R., Fernandez-Golbano, I. M., Plemel, R., Merz, A. J., Geli, M. I., Traub, L. M. and Lemmon, S. K. (2009). Clathrin functions in the absence of the terminal domain binding site for adaptor-associated clathrin-box motifs. Mol Biol Cell 20, 3401–13.

Drake, M. T. and Traub, L. M. (2001). Interaction of two structurally distinct sequence types with the clathrin terminal domain beta-propeller. J Biol Chem 276, 28700–9.

Dutta, D. and Donaldson, J. G. (2012). Search for inhibitors of endocytosis: Intended specificity and unintended consequences. Cellular logistics 2, 203–208.

Dutta, D., Williamson, C. D., Cole, N. B. and Donaldson, J. G. (2012). Pitstop 2 is a potent inhibitor of clathrin-independent endocytosis. PLoS One 7, e45799.

Edeling, M. A., Mishra, S. K., Keyel, P. A., Steinhauser, A. L., Collins, B. M., Roth, R., Heuser, J. E., Owen, D. J. and Traub, L. M. (2006). Molecular switches involving the AP-2 beta2 appendage regulate endocytic cargo selection and clathrin coat assembly. Dev Cell 10, 329–42.

Goodman, O. B., Jr., Krupnick, J. G., Gurevich, V. V., Benovic, J. L. and Keen, J. H. (1997). Arrestin/clathrin interaction. Localization of the arrestin binding locus to the clathrin terminal domain. J Biol Chem 272, 15017–22.

Heuser, J. E. and Anderson, R. G. (1989). Hypertonic media inhibit receptor-mediated endocytosis by blocking clathrin-coated pit formation. J Cell Biol 108, 389–400.

Hood, F. E., Williams, S. J., Burgess, S. G., Richards, M. W., Roth, D., Straube, A., Pfuhl, M., Bayliss, R. and Royle, S. J. (2013). Coordination of adjacent domains mediates TACC3-ch-TOG-clathrin assembly and mitotic spindle binding. J Cell Biol 202, 463–78.

Kang, D. S., Kern, R. C., Puthenveedu, M. A., von Zastrow, M., Williams, J. C. and Benovic, J. L. (2009). Structure of an arrestin2-clathrin complex reveals a novel clathrin binding domain that modulates receptor trafficking. J Biol Chem 284, 29860–72.

Knuehl, C., Chen, C. Y., Manalo, V., Hwang, P. K., Ota, N. and Brodsky, F. M. (2006). Novel binding sites on clathrin and adaptors regulate distinct aspects of coat assembly. Traffic 7, 1688–700.

Krupnick, J. G., Goodman, O. B., Jr., Keen, J. H. and Benovic, J. L. (1997). Arrestin/clathrin interaction. Localization of the clathrin binding domain of nonvisual arrestins to the carboxy terminus. J Biol Chem 272, 15011–6.

Larkin, J. M., Brown, M. S., Goldstein, J. L. and Anderson, R. G. (1983). Depletion of intracellular potassium arrests coated pit formation and receptor-mediated endocytosis in fibroblasts. Cell 33, 273–85.

Lemmon, S. K. and Traub, L. M. (2012). Getting in Touch with the Clathrin Terminal Domain. Traffic.

Macgregor, K. A., Robertson, M. J., Young, K. A., von Kleist, L., Stahlschmidt, W., Whiting, A., Chau, N., Robinson, P. J., Haucke, V. and McCluskey, A. (2014). Development of 1,8-naphthalimides as clathrin inhibitors. J Med Chem 57, 131–43.

McMahon, H. T. and Boucrot, E. (2011). Molecular mechanism and physiological functions of clathrin-mediated endocytosis. Nat Rev Mol Cell Biol 12, 517–33.

Miele, A. E., Watson, P. J., Evans, P. R., Traub, L. M. and Owen, D. J. (2004). Two distinct interaction motifs in amphiphysin bind two independent sites on the clathrin terminal domain beta-propeller. Nat Struct Mol Biol 11, 242–8.

Motley, A., Bright, N. A., Seaman, M. N. and Robinson, M. S. (2003). Clathrin-mediated endocytosis in AP-2-depleted cells. J Cell Biol 162, 909–18.

Smith, C. M., Haucke, V., McCluskey, A., Robinson, P. J. and Chircop, M. (2013). Inhibition of clathrin by pitstop 2 activates the spindle assembly checkpoint and induces cell death in dividing HeLa cancer cells. Molecular cancer 12, 4.

Stahlschmidt, W., Robertson, M. J., Robinson, P. J., McCluskey, A. and Haucke, V. (2014). Clathrin terminal domain-ligand interactions regulate sorting of mannose 6-phosphate receptors mediated by AP-1 and GGA adaptors. J Biol Chem.

ter Haar, E., Harrison, S. C. and Kirchhausen, T. (2000). Peptide-in-groove interactions link target proteins to the beta-propeller of clathrin. Proc Natl Acad Sci USA 97, 1096–100.

ter Haar, E., Musacchio, A., Harrison, S. C. and Kirchhausen, T. (1998). Atomic structure of clathrin: a beta propeller terminal domain joins an alpha zigzag linker. Cell 95, 563–73.

von Kleist, L. and Haucke, V. (2012). At the crossroads of chemistry and cell biology: inhibiting membrane traffic by small molecules. Traffic 13, 495–504.

von Kleist, L., Stahlschmidt, W., Bulut, H., Gromova, K., Puchkov, D., Robertson, M. J., MacGregor, K. A., Tomilin, N., Pechstein, A., Chau, N. et al. (2011). Role of the clathrin terminal domain in regulating coated pit dynamics revealed by small molecule inhibition. Cell 146, 471–84.

Willox, A. K. and Royle, S. J. (2012). Functional analysis of interaction sites on the N-terminal domain of clathrin heavy chain. Traffic 13, 70–81.

